# Mitochondria-adaptor TRAK1 promotes kinesin-1 driven transport in crowded environments

**DOI:** 10.1101/2020.01.22.915066

**Authors:** Verena Henrichs, Lenka Grycova, Cyril Barinka, Zuzana Nahacka, Jiri Neuzil, Stefan Diez, Jakub Rohlena, Marcus Braun, Zdenek Lansky

## Abstract

Intracellular trafficking of organelles, driven by kinesin-1 stepping along microtubules, underpins essential processes including neuronal activity. In absence of other proteins on the microtubule surface, kinesin-1 performs micron-long runs. Under protein crowding conditions, however, kinesin-1 motility is drastically impeded. It is thus unclear how kinesin-1 acts as an efficient transporter in crowded intracellular environments. Here, we demonstrate that TRAK1 (Milton), an adaptor protein essential for mitochondrial trafficking, activates kinesin-1 and increases its robustness of stepping in protein crowding conditions. Interaction with TRAK1 i) facilitated kinesin-1 navigation around obstacles, ii) increased the probability of kinesin-1 passing through cohesive envelopes of tau and iii) increased the run length of kinesin-1 in cell lysate. We explain the enhanced motility by the observed direct interaction of TRAK1 with microtubules, providing an additional anchor for the kinesin-1-TRAK1 complex. We propose adaptor-mediated tethering as a mechanism regulating kinesin-1 motility in various cellular environments.

## Introduction

In eukaryotic cells, microtubules constitute a major part of the cytoskeleton and provide a multi-functional scaffold crucial for intracellular long-range transport. Microtubule-based transport is particularly important in neurons, where it enables an efficient distribution of cargo, such as mitochondria, along elongated axons to distal regions of the cell. Active mitochondrial transport is equally essential for the redistribution of mitochondria during mitosis ^1, 2^ and trafficking of mitochondria between cells through tunnelling nanotubes (Ahmad et al., 2014; Dong et al., 2017; Rustom et al., 2004). Mitochondrial trafficking between stroma and cancer cells is relevant in tumour initiation and progression ^3, 8, 9^ and dysfunctions in the distribution of mitochondria are connected with neurodegenerative diseases, including Alzheimer’s disease, Huntington’s disease or amyotrophic lateral sclerosis ^10–15^.

Trafficking of cargo, including mitochondria, is driven by molecular motors such as kinesin-1 ^16^. By hydrolysing ATP in its N-terminal motor domains, kinesin-1 heavy chain (further referred to as kinesin-1) moves in steps towards the plus-end of microtubules. Thus, kinesin-1 drives anterograde transport ^17–19^ of mitochondria, which are transported in cells in bursts of motion, covering distances of tens of micrometresi) ^20^. Kinesin-1, in absence of other proteins on the microtubule surface *in vitro*, is processive, meaning that it can perform more than 100 consecutive steps towards the microtubule plus-end, covering hundreds of nanometres before dissociating from the microtubule ^21^. The coupling of multiple molecular motors to a single cargo further increases the cargo processivity, as shown *in vitro* in the absence of other proteins on the microtubule surface ^22–25^. Indeed, various cellular cargoes are transported by multiple molecular motors (Kural et al., 2005; Levi et al., 2006; Welte et al., 1998). In cells, however, microtubules are heavily decorated by a large variety of proteins, crowding the microtubule surface ^29^, which strongly impedes kinesin-1-driven transport through a drastic reduction of kinesin-1 processivity (Telley et al., 2009). Mechanisms additional to coupling of multiple motors are thus likely at play to overcome the hindering effect of crowding and to enable robust long-range kinesin-1-driven transport in cells.

Binding of cargo, such as mitochondria, is mediated by adaptor proteins interacting with the C-terminal tail-domain of kinesin-1. In the absence of cargo, kinesin-1 is auto-inhibited via an interaction of the cargo-binding domain with the motor domain ^32–34^. Mitochondria as cargo are physically linked to kinesin-1 by the adaptor proteins Miro and Milton ^35–40^. Two mammalian Milton homologues are known, TRAK1 and TRAK2, (trafficking kinesin protein 1 and 2, respectively), with TRAK1 being the preferred binding partner of kinesin-1 ^41, 42^. The binding of TRAK1 to kinesin-1 is mediated by a direct association of the coiled-coil domain of TRAK1 with the cargo-binding site in the tail-domain of kinesin-1 ^36, 39, 41–44^. Since a direct interaction of a protein with a molecular motor is likely to alter the motor’s transport properties and cargo-binding to the auto-inhibitory regions might activate motors ^32, 33, 45^, we investigated how the adaptor protein TRAK1 modulates the motility of kinesin-1.

Here we report that TRAK1 activates kinesin-1 and promotes long-range transport on densely crowded microtubules by increasing the molecular motors’ processivity. We explain these observations by TRAK1-mediated anchoring of kinesin-1 to the microtubule surface, and we propose auxiliary anchoring, mediated by adaptor proteins, as a mechanism regulating kinesin-1 transport, which is particularly supportive in crowded environments.

## Results

### TRAK1 diffuses along microtubules and activates human kinesin-1 KIF5B

Full-length KIF5B (herein referred to as KIF5B; Figure S1a) is a kinesin-1 heavy chain encoded in the human genome and it is auto-inhibited in the absence of a cargo ^32^, unable to walk processively along microtubules. In order to study the interaction of KIF5B with TRAK1, we immobilized microtubules onto a coverslip surface and added mCherry-labelled full-length TRAK1 (mCherry-TRAK1; Figure S1a, S1b, left panel) and/or GFP-labelled full-length KIF5B (KIF5B-GFP; Figure S1a). To visualize the molecular interactions with the microtubules we used total internal reflection fluorescence (TIRF) microscopy with single molecule resolution (Methods). In the absence of KIF5B-GFP, we observed mCherry-TRAK1 diffusing along microtubules (Figure 1a), demonstrating that TRAK1, rather unexpectedly, contains a microtubule-binding domain. Imaging of KIF5B-GFP in the absence of TRAK1 showed that it indeed was essentially auto-inhibited, unable to bind to microtubules (Figure S1c). Consistent with previously published data ^32, 46, 47^, we observed occasional brief diffusion of KIF5B-GFP molecules along microtubules and sporadic processive runs (Fig. 1c). By contrast, when imaging KIF5B-GFP in the presence of mCherry-TRAK1, we observed, at the same concentration of KIF5B-GFP, an order of magnitude increase in the number of molecules performing processive runs (Figure 1b, 1c, n = 160 molecules). The fluorescence signal of KIF5B-GFP and mCherry-TRAK1 colocalized in about 40% of the processive runs (Figure 1b, 1c), demonstrating that TRAK1 and KIF5B formed a processive transport complex. In about 10% of the cases a processive run was detected only in the mCherry channel and in about 50% of the cases only in the GFP channel (Figure 1c). We assume that these runs were performed by the KIF5B-TRAK1 transport complex with one of the fluorescent proteins photo-bleached as neither of the proteins individually was capable of robust processive movement. These results show that the adaptor protein TRAK1 interacts directly with microtubules and activates processive motion of KIF5B.

**Figure 1.**
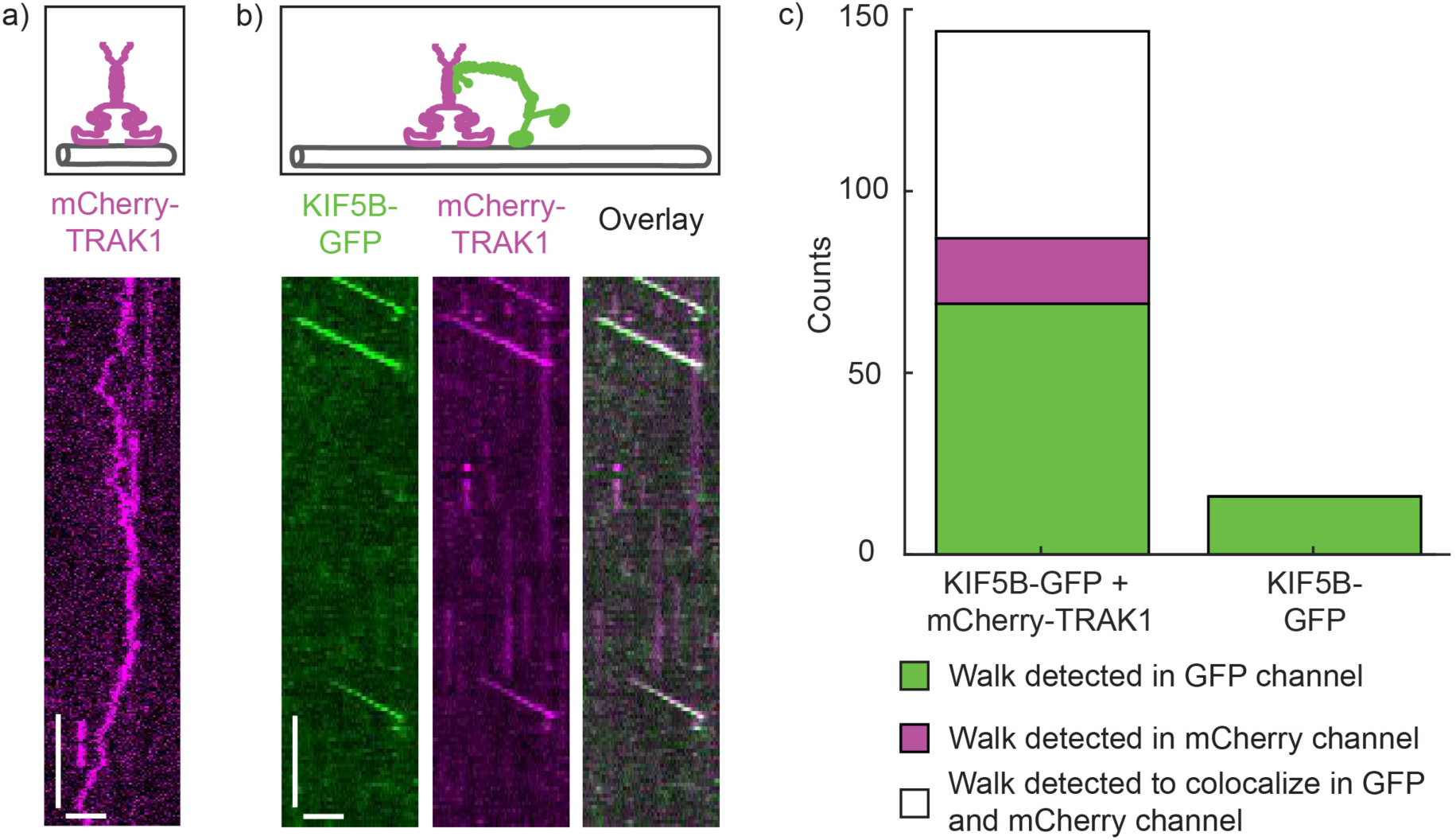
TRAK1 activates KIF5B. Schematic illustrations of the experimental set-up and kymographs of **a)** single mCherry-TRAK1 molecules (magenta) moving diffusively along a microtubule and **b)** mCherry-TRAK1 molecules (magenta) colocalizing with KIF5B-GFP molecules (green), which move processively along microtubules, showing that mCherry-TRAK1 activates KIF5B-GFP. Scale bars: 2 µm, 10 s. **c)** Number of processively walking KIF5B-GFP molecules in the presence (left) and absence of mCherry-TRAK1 (right) detected in the GFP channel (green), mCherry channel (magenta) or colocalizing in both channels (white).

### KIF5B transports two TRAK1 dimers

To be able to directly compare the influence of TRAK1 on the KIF5B motility, we generated a constitutively active KIF5B construct by removing the KIF5B inhibitory domain (amino acids 906-963, KIF5BΔ; Figure S1a). Analogously to the experiment above, we first visualized GFP-labelled KIF5BΔ (KIF5BΔ-GFP; Figure S1a) interacting with microtubules, confirming that it moved processively (Figure 2a). Next, we imaged the interaction of KIF5BΔ-GFP with mCherry-TRAK1 on microtubules. Similar to full-length KIF5B-GFP, we observed colocalization of KIF5BΔ-GFP and mCherry-TRAK1 during processive runs (Figure 2b), showing that the removal of the inhibitory sequence of KIF5B did not disrupt the KIF5B interaction with TRAK1. In all following experiments we thus used this constitutively active KIF5B construct.

**Figure 2.**
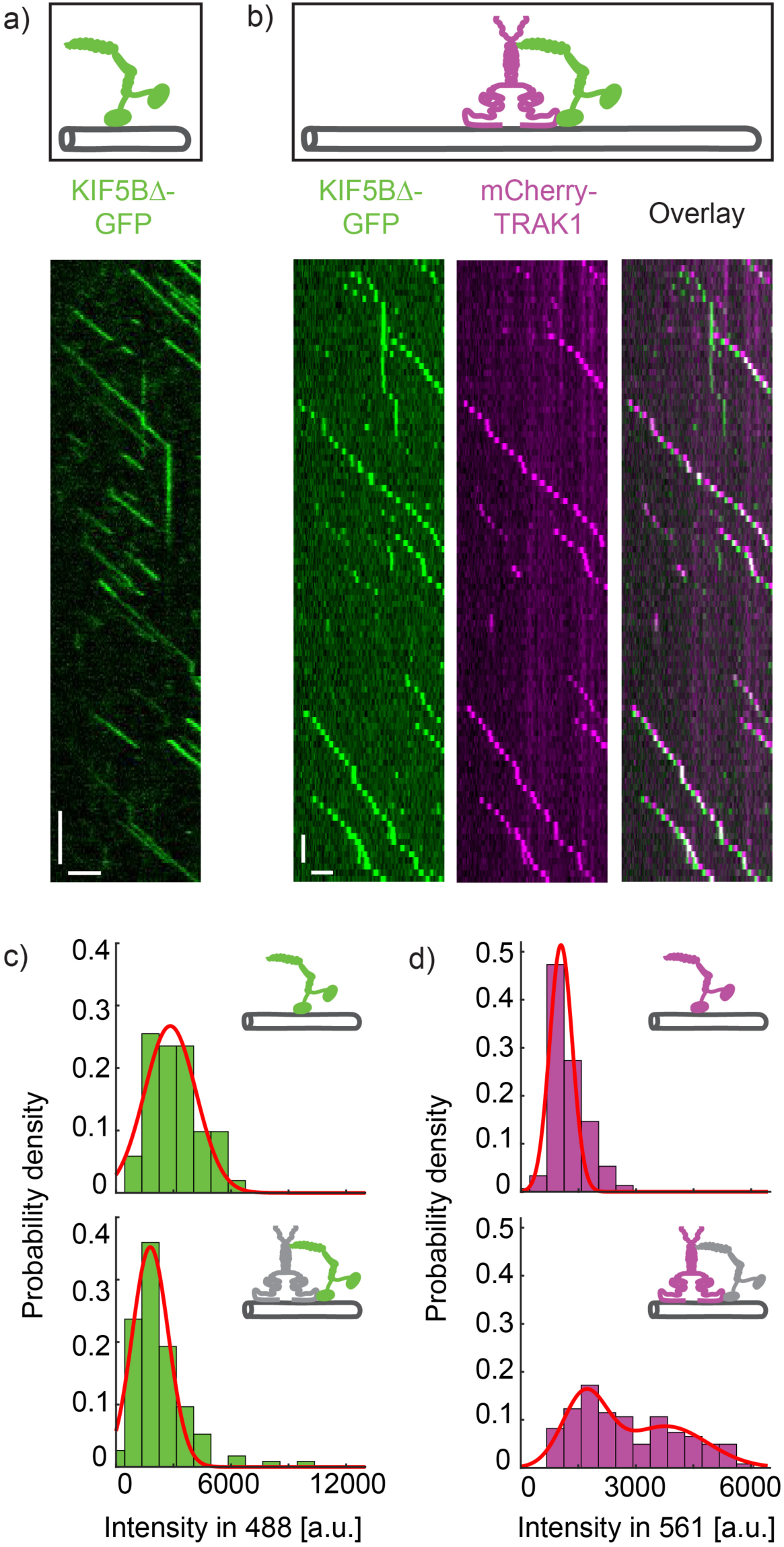
KIF5B transports two TRAK1 dimers. Schematic illustrations of the experimental set-up and kymographs of **a)** KIF5BΔ-GFP (green) and **b)** KIF5BΔ-GFP (green) colocalizing with mCherry-TRAK1 (magenta), walking processively along microtubules. Scale bars: 2 µm, 5 s **c)** Histograms of the integrated fluorescence intensity and Gaussian fits (red) of the GFP-signal of KIF5BΔ-GFP in the absence (top) and presence (bottom) of mCherry-TRAK1. Both distributions comprise one peak only, indicating that one KIF5BΔ dimer is involved in the transport complex. See also Figure S2a. **d)** Histograms of the integrated fluorescence intensity and Gaussian fits (red) of the mCherry-signal of KIF5BΔ-mCherry (top) and of mCherry-TRAK1 in complex with KIF5BΔ-GFP (bottom), showing two peaks in the latter case, indicating that two TRAK1 dimers can be transported by KIF5B. See also Figure S2b.

During the processive movement of KIF5BΔ-GFP-mCherry-TRAK1 complexes, we observed multiple bleaching steps of the fluorescence signal of KIF5BΔ-GFP and mCherry-TRAK1 (Figure S2), suggesting possible multimerization states of the two proteins. We thus sought to identify the number of KIF5BΔ-GFP and mCherry-TRAK1 molecules involved in the processive transport complexes. We first compared the fluorescence intensities of processively moving KIF5BΔ-GFP (Methods), which is known to be in a dimeric form ^30, 48^, with the signal of KIF5BΔ-GFP in complex with mCherry-TRAK1. The probability density distribution of the background-corrected fluorescence intensity of KIF5BΔ-GFP in the absence of mCherry-TRAK1 exhibited a single population, which we interpret as the signal of two GFP molecules (Figure 2c top, n = 51 molecules, Methods). As the probability density distribution of the background-corrected fluorescence intensity of KIF5BΔ-GFP in the presence of mCherry-TRAK1 exhibited also a single population (Figure 2c bottom, n = 114 molecules), similar to KIF5BΔ-GFP alone, we conclude that the transport complex is formed by a single KIF5BΔ dimer.

We next analogously estimated the number of mCherry-TRAK1 molecules in a processive transport complex. As a standard for the fluorescence intensity of an mCherry-labelled dimer, we evaluated the background-corrected fluorescence intensity of mCherry-labelled KIF5BΔ (KIF5BΔ-mCherry; Figure 2d top, n = 150 molecules; Figure S1a). Assessing the fluorescent intensities of mCherry-TRAK1 in the presence of KIF5BΔ-GFP resulted in two populations in the probability density distribution (Figure 2d bottom, n = 122 molecules). In agreement with previously published data demonstrating TRAK1 dimerization ^40^, we found that one population exhibits intensities comparable to the intensities observed for a KIF5BΔ-mCherry dimer. A second population was located at approximately two times higher values, consistent with intensities expected for four mCherry-fluorophores of two mCherry-TRAK1 dimers. These experiments demonstrate that processive KIF5B-TRAK1 transport complexes comprise one KIF5B dimer and up to two TRAK1 dimers.

### TRAK1 increases the processivity of KIF5B

In the experiments described above, we noticed that KIF5BΔ in the presence of TRAK1 performs longer runs than in the absence of TRAK1. To quantify the effect of TRAK1 on the KIF5BΔ motility, we determined the velocity, the interaction time with microtubules and the lengths of individual processive runs of KIF5BΔ-GFP and of KIF5BΔ-GFP-mCherry-TRAK1 complexes (exemplarily shown in Figure 3a, 3b). In the latter condition, to avoid accounting for KIF5BΔ-GFP molecules that are not in complex with mCherry-TRAK1, we analysed only traces of mCherry-TRAK1 (which by itself never performs processive runs; Figure 1a). Strikingly, we observed an increase of the median run length of KIF5BΔ-GFP in the presence of mCherry-TRAK1 by several fold from 1.75 µm (95% confidence interval, CI_95_ (1.55, 2.03) µm, n = 499 molecules) to 5.70 µm (CI_95_ (3.78, 6.87) µm, n = 222 molecules) (Figure 3e). Similarly, the median interaction time of KIF5BΔ-GFP with microtubules increased by an order of magnitude from 1.84 s (CI_95_ (1.64, 2.05) s, n = 534 molecules) in the absence of mCherry-TRAK1 to 12.20 s (CI_95_ (8.54, 15.86) s, n = 221 molecules) in its presence (Figure 3f). Simultaneously, the velocity of KIF5BΔ-GFP decreased in the presence of mCherry-TRAK1 from 918 ± 171 nm/s (mean ± standard deviation, n = 534 molecules) to 604 ± 263 nm/s (mean ± standard deviation, n = 499 molecules; Figure 3d, Methods). In addition to the reduction of velocity, we observed a higher occurrence of transient pauses in the presence of mCherry-TRAK1 (Figure S3a).

**Figure 3.**
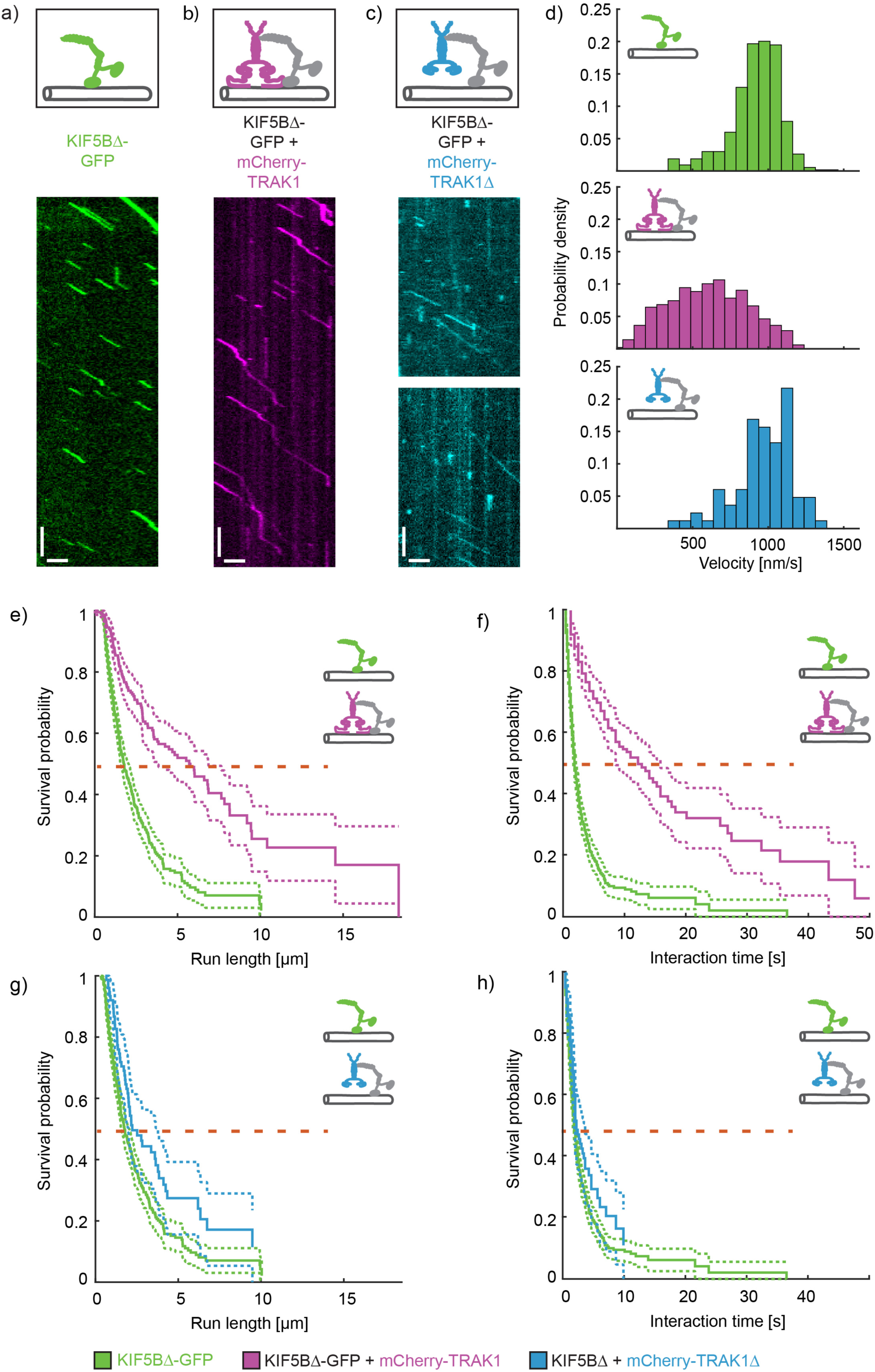
TRAK1 increases the processivity of KIF5B. Schematic illustrations of the experimental set-up and kymographs showing processive movement of **a)** KIF5BΔ-GFP (green), **b)** mCherry-TRAK1 (magenta) in the presence of KIF5BΔ-GFP (not imaged) and **c)** mCherry-TRAK1Δ (cyan) in the presence of KIF5BΔ-GFP (not imaged). Scale bars: 2 µm, 5 s. See also Figure S3b. **d)** Histograms of the velocities of KIF5BΔ-GFP (green, top), KIF5BΔ-GFP in the presence of mCherry-TRAK1 (magenta, middle) and KIF5BΔ-GFP in the presence of mCherry-TRAK1Δ (cyan, bottom). The velocity of KIF5BΔ-GFP decreased only in the presence of mCherry-TRAK1 but not in presence of mCherry-TRAK1Δ. See also Figure S3a. **e), f)** Survival probability (Kaplan-Meier estimation) of the run length and interaction time, respectively, of KIF5BΔ-GFP in the absence (green) and presence of mCherry-TRAK1 (magenta). In the presence of mCherry-TRAK1, the median run length and median interaction time of KIF5BΔ-GFP increased. **g), h)** Survival probability showing that mCherry-TRAK1Δ (cyan) hardly affects the run length and interaction time of KIF5BΔ-GFP (green). The dashed horizontal lines highlight the median run lengths and interaction times, estimated as the shortest survival time, for which the survival probability is equal to or less than 0.5.

We hypothesized that the increase in the run length and interaction time of KIF5BΔ in the presence of TRAK1 is caused by the microtubule-binding domain of TRAK1, which may anchor KIF5BΔ to the microtubule and thus decrease the probability of KIF5BΔ dissociating from the microtubule. The observed simultaneous decrease of the velocity of the KIF5BΔ-TRAK1 complex is in agreement with this notion, as a diffusible TRAK1 interaction with the microtubule would likely generate an additional frictional drag, slowing down the molecular motor. The C-terminal region of TRAK1 exhibits a high isoelectric point of 10, and is thus likely to interact electrostatically with the negatively charged microtubule. To examine if this region contributes to the increased processivity of KIF5BΔ, we deleted the C-terminus of mCherry-labelled TRAK1 (mCherry-TRAK1Δ; Figure S1a, S1b, right panel), which prevented its interaction with microtubules (Figure S3b) but did not compromise its binding to KIF5BΔ-GFP (Figure 3c). We then evaluated the run length and interaction time of KIF5BΔ-GFP in complex with mCherry-TRAK1Δ. Consistent with the hypothesis that TRAK1Δ does not anchor KIF5BΔ to microtubules, mCherry-TRAK1Δ barely affected the velocity of KIF5BΔ-GFP (972 ± 196 nm/s, mean ± standard deviation, n = 82 molecules; Figure 3d), its median run length (2.22 µm, CI_95_ (1.99, 3.78) µm, n = 82 molecules) or interaction time (2.22 s, CI_95_ (1.82, 3.63) s, n = 82 molecules) (Figure 3g, 3h). We thus conclude that the microtubule-binding domain located at the C-terminus of TRAK1 provides an additional anchor point for KIF5B, which decreases the probability of the KIF5B-TRAK1 transport complex to dissociate from the microtubule and increases the run length as well as the interaction time of the complex, albeit at the cost of adding an additional frictional drag on KIF5B, slowing the molecular motor down.

### TRAK1 increases the processivity of KIF5B in crowded environments

The microtubule surface in cells is crowded with numerous microtubule-associated proteins hindering the motion of molecular motors. Crowding of the microtubule surface results in a drastic shortening of the run length of kinesin-1 ^30, 31, 49, 50^. Since we observed that TRAK1 increased the processivity of KIF5B, we wondered whether TRAK1 can extend the movement range of kinesin-1 in crowded conditions. We thus repeated the experiments described above in three different conditions of crowding.

i) First, we used a GFP-labelled rigor-binding truncated kinesin-1 T93N mutant (Schneider et al., 2015) (Methods, Figure 4a-c and S1a). This mutant is unable to hydrolyse ATP ^51^ and therefore binds with a high affinity to microtubules forming stationary roadblocks that perfectly mask the KIF5B binding sites. As described before ^31^, the median run length of KIF5BΔ-mCherry decreased in the presence of static roadblocks by about 50% from 1.75 µm (CI_95_ (1.55, 2.03) µm, n = 499 molecules, same dataset as in Figure 3d-h) to 0.86 µm (CI_95_ (0.79, 0.98) µm, n = 547 molecules) (Figure 4b). Strikingly, addition of mCherry-TRAK1 increased the median run length of KIF5BΔ-mCherry in the presence of the roadblocks about two-fold, to 1.12 µm (CI_95_ (1.06, 1.29) µm, n = 1066 molecules; Figure 4b). Consistently, the median interaction time of KIF5BΔ-mCherry decreased in the presence of the roadblocks by about 30% from 1.84 s (CI_95_ (1.64, 2.05) s, n = 534 molecules, same dataset as in Figure 3d-h) to 1.40 s (CI_95_ (1.20, 1.43) s, n = 547 molecules) (Figure 4c). The presence of mCherry-TRAK1 restored the median interaction time of KIF5BΔ-mCherry in the presence of roadblocks to 1.84 s (CI_95_ (1.80, 2.20) s, n = 1066 molecules; Figure 4c). These data demonstrate that TRAK1 promotes KIF5BΔ processivity even when the exact binding sites for the motor-domain are occupied.

**Figure 4.**
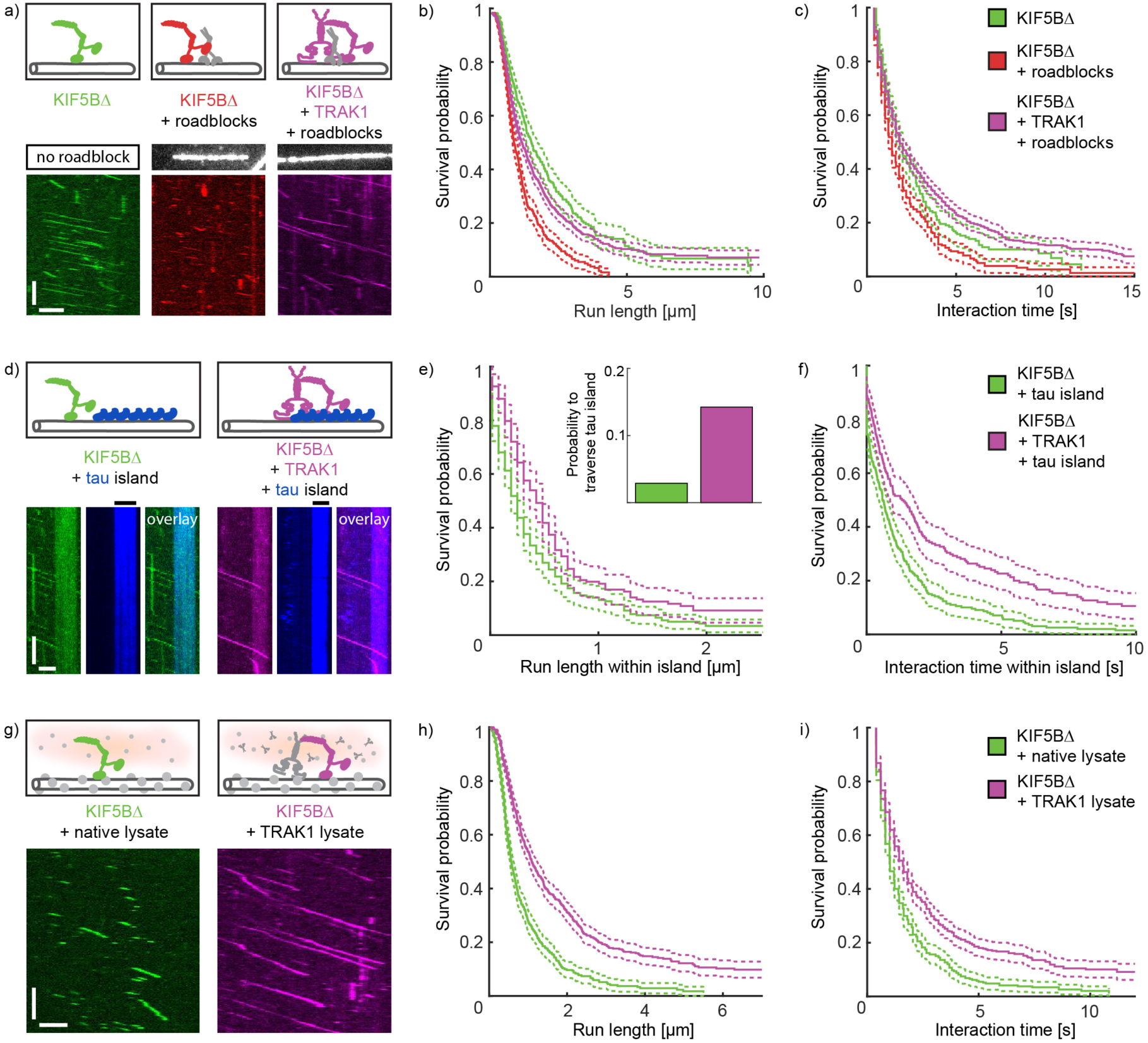
TRAK1 increases the processivity of KIF5B in crowded environments. **a)** Schematic illustrations of the experimental set-up and kymographs showing processive movement of KIF5BΔ-GFP (left, green), KIF5BΔ-GFP in the presence of roadblocks (middle, red) and KIF5BΔ-GFP in the presence of mCherry-TRAK1 and roadblocks (right, magenta). **b), c)** Survival probability of the run length and interaction time of KIF5BΔ-GFP (green), KIF5BΔ-GFP in the presence of roadblocks (red) and KIF5BΔ-GFP in the presence of mCherry-TRAK1 and roadblocks (magenta). The median run length and median interaction time decreased for KIF5BΔ-GFP in the presence of roadblocks but increased in presence of mCherry-TRAK1 despite the presence of roadblocks. The control of KIF5BΔ-GFP comprises the same datasets as in Figure 3d-h. **d)** Schematic illustrations of the experimental set-up and kymographs showing processive movement of KIF5BΔ-GFP in the absence (left panel, green) and presence of mCherry-TRAK1 (right panel, magenta) on tau-mCherry island-decorated microtubules (blue). The positions of the tau-mCherry islands are denoted by black horizontal lines. KIF5BΔ-GFP dissociated from the microtubule at the edges of tau-mCherry islands (left panel) but penetrated further into the tau-mCherry islands in the presence of mCherry-TRAK1 (right panel). **e), f)** Survival probability plots showing the increased median run length and median interaction time of KIF5BΔ-GFP within tau-mCherry islands in the presence of mCherry-TRAK1 (magenta) in comparison to KIF5BΔ-GFP in the absence of mCherry-TRAK1 (green). Inset: KIF5BΔ-GFP in the presence of mCherry-TRAK1 (magenta) showed a higher probability to traverse an entire tau-mCherry island than KIF5BΔ-GFP alone (green). For a distribution of island lengths see Figure S4a, S4b. **g)** Schematic illustrations of the experimental set-up and kymographs showing processive movement of KIF5BΔ-GFP in native cell lysate (left, green) and in TRAK1-overexpressing cell lysate (right, magenta). **h), i)** Survival probability plots showing the increased median run length and median interaction time of KIF5BΔ-GFP in TRAK1 lysate (magenta) in comparison to native lysate (green). See also Fig. S4c, S4d. Scale bars: 2 µm, 10 s.

ii) To test if TRAK1 anchoring promotes KIF5BΔ-GFP processivity also in the presence of intrinsically disordered, microtubule-bound, neuronal proteins that regulate axonal transport, we next used the unstructured, microtubule-associated mCherry-labelled protein tau (tau-mCherry). Tau forms cohesive islands of high density on the microtubule surface ^52^, preventing kinesin-1 from walking into regions of microtubules covered by these islands ^53^ (Figure 4d). When KIF5BΔ-GFP encountered a tau-mCherry island, about 35% of the molecules detached directly at the boundary of the island. The remaining molecules penetrated the islands to reach a median run length within the island of 0.17 µm (CI_95_ (0.17, 0.23) µm, n = 242 molecules; Figure 4e), before detachment. By contrast, complexes of KIF5BΔ-GFP and mCherry-TRAK1 detached in about 17% of the cases while the median run length within tau-mCherry islands, when compared to KIF5BΔ-GFP alone, increased about threefold to 0.41 µm (CI_95_ (0.29, 0.46) µm, n = 203 molecules; Figure 4e). This increase in the processivity within the tau-mCherry islands resulted in an about fivefold increased probability of KIF5BΔ-GFP to completely traverse a tau-mCherry island when in complex with mCherry-TRAK1 (Figure 4e, inset; Figure S4a, S4b). Consistently, the median interaction time of KIF5BΔ-GFP within tau-mCherry islands increased in the presence of mCherry-TRAK1 from 0.59 s (CI_95_ (0.46, 0.77) s, n = 242 molecules) to 1.31 s (CI_95_ (0.87, 1.78) s, n = 203 molecules) (Figure 4f). These results demonstrate that TRAK1-mediated anchoring increases the efficiency of KIF5B-driven transport within regions of microtubules coated by cohesive envelopes of intrinsically-disordered microtubule-associated neuronal proteins.

iii) Having examined the role of TRAK1-mediated anchoring on KIF5B stepping behaviour in the presence of distinct microtubule-binding proteins, we next moved to the more complex milieu of whole cell lysates containing a number of various microtubule-associated proteins. To this end, we lysed native HEK-293/T17 cells, HEK-293/T17 cells overexpressing mCherry-TRAK1 (further denoted as TRAK1 lysate) and HEK-293/T17 cells overexpressing the purification tag only (further denoted as Halo lysate, Methods). We supplemented these lysates with KIF5BΔ-GFP to visualize its motion along microtubules (Figure 4g). The median run length of KIF5BΔ-GFP in the native cell lysate was 0. 57 µm (CI_95_ (0.53, 0.63) µm, n = 535 molecules), and thereby, as previously reported ^30^, much shorter than on bare microtubules (Figure 4a, 4b, 4h). Strikingly, in TRAK1 lysate, the median run length increased about two-fold to 1.09 µm (CI_95_ (0.99, 1.23), n = 715 molecules) (Figure 4h). Consistently, the median interaction time of KIF5BΔ-GFP increased by 60% from 1.02 s (CI_95_ (1.00, 1.20) s, n = 535 molecules) in native cell lysate to 1.60 s (CI_95_ (1.43, 1.80) s, n = 715 molecules) in TRAK1 lysate (Figure 4i). This increase in the run length or interaction time was not observed in Halo lysate overexpressing the purification tag (Figure S4c, S4d, run length 0.63 µm CI_95_ (0.58, 0.69) µm, interaction time 1.00 s CI_95_ (0.82, 1.02) s, n = 559 molecules), demonstrating that TRAK1 enhances the KIF5B processivity on highly crowded microtubules under native conditions in cell extract. Taken together, our results show that TRAK1 promotes long range KIF5BΔ-based transport on crowded microtubules.

### TRAK1 enables mitochondrial transport in vitro

Finally, we asked if the KIF5B-TRAK1 transport complex characterized above can transport mitochondria. We thus isolated mitochondria expressing mitochondria-targeted eGFP (denoted as mitochondria-GFP) from 4T1 cells using a mild isolation protocol (Methods). We verified by mass spectrometry that these mitochondria did not contain TRAK1 or KIF5B, but the transmembrane protein Miro (see supplementary file *Mass Spectrometry of mitochondria*), providing the linkage between mitochondria and the transport complex ^38^. We then combined mitochondria-GFP with recombinant mCherry-TRAK1 and unlabelled KIF5BΔ, and added this mixture to surface-attached microtubules (Methods). We observed that mitochondria-GFP colocalized on microtubules with mCherry-TRAK1 and moved processively along the microtubules (Figure 5a). We also observed processive movement of mCherry-TRAK1, presumably in complex with unlabelled KIF5BΔ. In control experiments, the absence of TRAK1 and/or KIF5BΔ prevented processive movement of mitochondria-GFP along microtubules. To compare the motility of single mitochondria-GFP with the motility of KIF5BΔ-GFP and of KIF5BΔ-GFP-mCherry-TRAK1 complexes, we analysed the respective velocities and the corresponding run lengths (Methods). As described above, single KIF5BΔ-GFP moved with the highest velocity of 918 ± 171 nm/s (mean ± standard deviation, n = 534 molecules), while the velocity of KIF5BΔ-GFP in the presence of mCherry-TRAK1 was reduced to 604 ± 263 nm/s (mean ± standard deviation, n = 230 molecules) (Figure 5b, top and middle, same dataset as in Figure 3d-h). Mitochondria-GFP moved with a velocity of 266 ± 148 nm/s (mean ± standard deviation, n = 72 mitochondria; Figure 5b bottom). Consistently, KIF5BΔ-GFP exhibited the shortest median run length of 1.75 µm (CI_95_ (1.55, 2.03) µm, n = 499 molecules), which was increased to 5.70 µm (CI_95_ (3.78, 6.86) µm, n = 222 molecules) in the presence of mCherry-TRAK1 (Figure 5c, same dataset as in Figure 3d-h). The median run length of mitochondria-GFP was determined to be longer than 20 µm (n = 72 mitochondria; Figure 5c; Methods). The low velocity and high run length of mitochondria-GFP suggests that several KIF5BΔ-TRAK1 transport complexes might be attached to a single mitochondrion. In summary, we show that supplementing isolated mitochondria with recombinant TRAK1 and KIF5BΔ enables processive mitochondrial motility along microtubules. Moreover, we show that mitochondria move slower but travel further than single KIF5BΔ molecules or KIF5BΔ-TRAK1 transport complexes.

**Figure 5.**
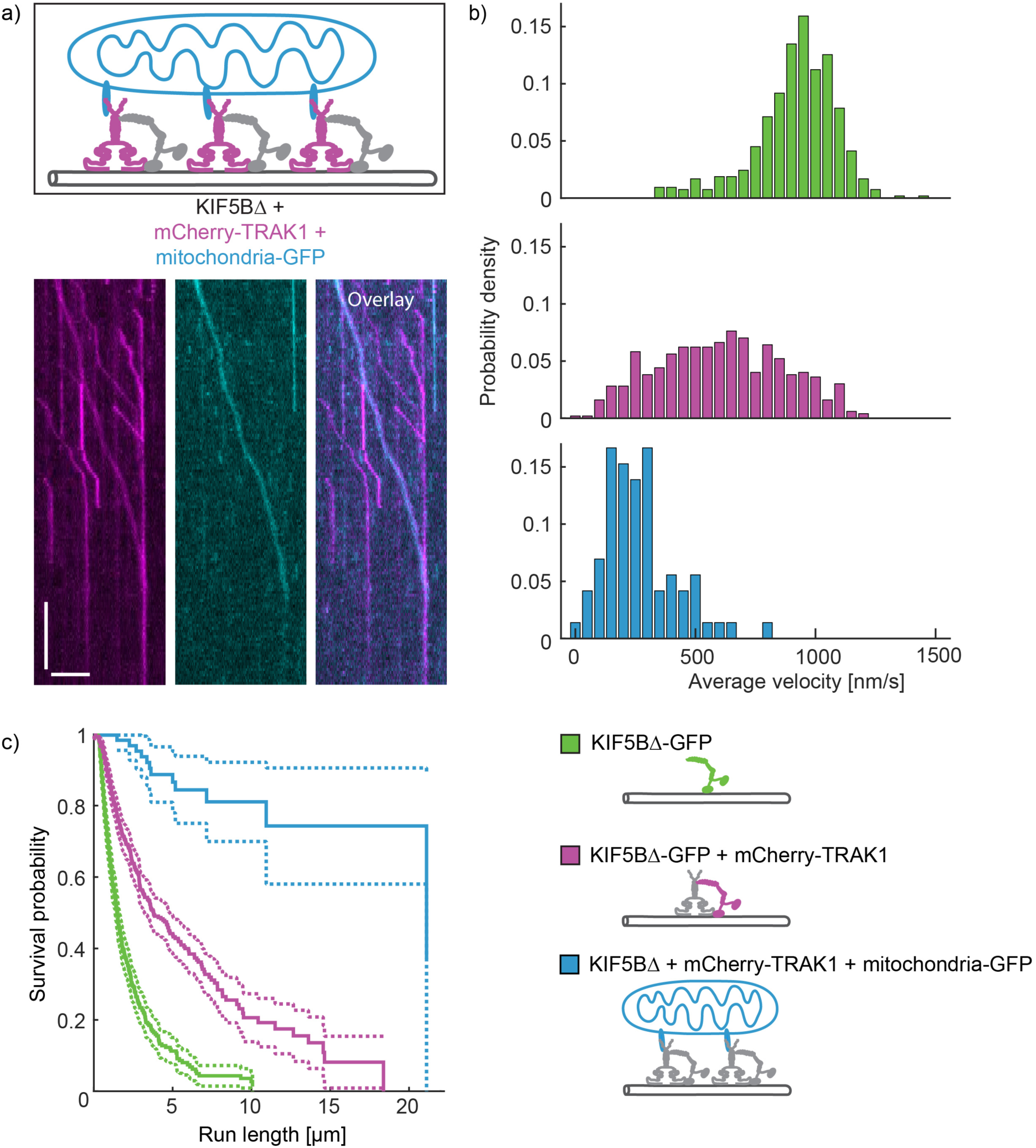
TRAK1 enables mitochondrial transport *in vitro*. **a)** Schematic illustration of the experimental set-up and kymographs showing mitochondria (cyan) being processively transported along microtubules by mCherry-TRAK1 (magenta), bound to unlabelled KIF5BΔ. **b)** Histograms showing the velocities of processive movement of KIF5BΔ-GFP (green), decreased in the presence of mCherry-TRAK1 (magenta) and further decreased for mitochondria-GFP transported by KIF5BΔ-mCherry-TRAK1 complexes (cyan). **c)** Survival probability plot showing an increased median run length of mitochondria-GFP transported by KIF5BΔ-mCherry-TRAK1 complexes (cyan) in comparison to KIF5BΔ-GFP alone (green) and to KIF5BΔ-GFP-mCherry-TRAK1 complexes (magenta). Scale bars 5 µm, 10 s. For the evaluation of the run length and velocity of KIF5BΔ-GFP in the absence and presence of mCherry-TRAK1, the same datasets as in Figure 2d-h were used. For a list of proteins present after the crude isolation of mitochondria refer to the supplementary file *Mass Spectrometry of mitochondria*.

## Discussion

We here demonstrate the innate function of TRAK1 to increase KIF5B processivity, i.e. the average number of consecutive steps that KIF5B performs, by directly tethering KIF5B to the microtubule. During the processive motion of dimeric kinesin-1 molecular motors, the motor domains alternatingly disengage and engage with the microtubule surface. At any one time, one motor domain is engaged to provide anchoring for the disengaged motor domain, which is searching for the next binding site on the microtubule. If the anchoring motor domain disengages before the next binding site is found, the molecular motor unbinds from the microtubule, which terminates its run. Therefore, a low detachment rate of the anchoring motor domain extends the time available for finding the next binding site and increases the processivity of the molecular motor ^54^. While binding of TRAK1 to kinesin-1 was reported previously ^36, 39, 41–44^, we here found that TRAK1, via its C-terminus, also directly interacts with the microtubule, diffusively tethering KIF5B to the microtubule surface. This tethering increases the processivity of the molecular motor, likely by extending the time available for the disengaged motor domain of KIF5B to search for the next free binding site on the microtubule (Figure 6). The ability of KIF5B to bind two TRAK1 dimers likely strengthens this effect by further reducing the probability of the complex to permanently dissociate from the microtubule. Analogously, the non-processive budding yeast kinesin-14 Kar3 becomes processive by forming a heterodimer with a non-motor microtubule-associated protein that anchors Kar3 to the microtubule ^55^. Likewise, the kinesin-3 KIF1A attains processivity as a single-headed monomer due to anchoring provided by the K-loop, a highly positively charged region internal to the KIF1A motor domain ^56, 57^. Increasing processivity by reciprocal anchoring has as well been reported for collective movement of assemblies containing increasing numbers of molecular motors ^22^. The anchoring role of TRAK1 could be further regulated e.g. by TRAK1 posttranslational modifications, which were shown to alter mitochondrial trafficking ^58^.

**Figure 6.**
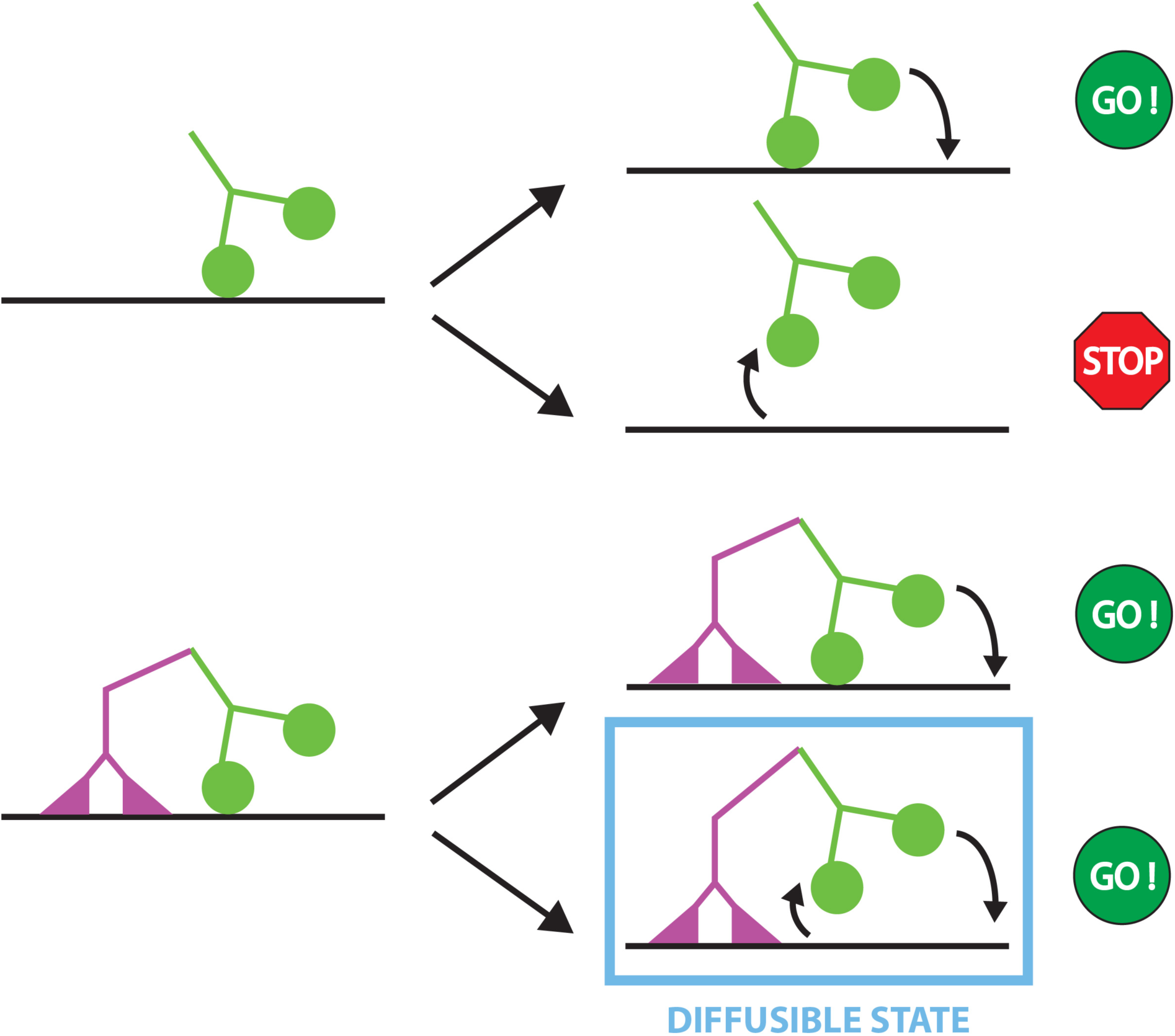
Schematic illustration of the TRAK1-mediated anchoring of KIF5B. Top: In the absence of TRAK1, KIF5B (green) can either continue its walk by rebinding the disengaged motor domain to the microtubule or dissociate from the microtubule when the engaged motor domain unbinds from the microtubule. Bottom: In presence of microtubule-bound TRAK1 (magenta), when both motor domains of KIF5B disengage from the microtubule, KIF5B remains tethered to the microtubule through a diffusive interaction of TRAK1 with the microtubule and thereby enables the rebinding of a motor domain of KIF5B to the microtubule. In this state, TRAK1 might facilitate navigation around obstacles by diffusion along the microtubule surface.

Microtubule surfaces in cells are crowded by numerous microtubule-associated proteins, which act as obstacles for processively moving molecular motors. When kinesin-1 encounters an obstacle, its run length decreases as kinesin-1 is more likely to dissociate from the microtubule ^30, 49^ than to continue its run by switching the protofilaments by side-stepping and circumventing the obstacle ^31^. *In vitro* single molecule experiments in cell lysate imitate the physiological state of microtubules crowded with microtubule associated proteins ^30, 59^, while rigor-binding kinesin-1 mutants provide static roadblocks masking the kinesin-1 binding sites ^31^. Our experiments showed that TRAK1 strongly increased the KIF5B processivity under these crowding conditions (schematically shown in Figure S5). We suggest that an additional anchoring of KIF5B by TRAK1 increases the time the molecular motor can pause in front of an obstacle without detaching from the microtubule. The extended time available increases the probability that either the obstacle vacates the occupied binding site or that the molecular motor side-steps to circumvent the obstacle.

The intrinsically disordered microtubule-associated protein tau presents a distinctive type of obstacle. In healthy neurons, the tau concentration increases from the cell body to the synapse ^60, 61^ whereas its distribution is reversed in neurodegenerating neurons ^62^. Tau is an important regulator of microtubule-based transport, by decreasing the run length of plus-end directed cargo transport ^63, 64^ it inhibits kinesin-dependent trafficking of mitochondria and thereby implicates Alzheimer’s disease ^65^. Knockdown of TRAK1 was demonstrated to result in neurodegeneration, which was suppressed by an additional knockdown of tau ^66^. *In vitro*, tau cooperatively forms cohesive islands on the microtubule surface, preventing kinesin-1 driven transport within the tau island-coated regions of microtubules ^52, 53^. Our results show that TRAK1 enables KIF5B entering regions covered with tau islands, increasing the probability of KIF5B to traverse tau islands and thus overriding the tau island-dependent blockade of kinesin-1-driven transport. Mechanistically, we propose that TRAK1-mediated anchoring to the microtubule increases the KIF5B waiting time in front of the tau island’s edge, promoting the probability of the KIF5B motor domain to displace the outmost of tau’s microtubule-binding repeats when the repeat transiently vacates its binding site. By this mechanism KIF5B can sequentially displace the tau microtubule-binding repeats, enabling motion through the island-covered microtubule region. Similarly, Kip3, a super-processive kinesin known to pause in the absence of an adjacent free binding site ^67^, traverses cohesive tau islands efficiently^53^. Other neuronal intrinsically disordered microtubule-associated proteins are likely to form similar cohesive islands on microtubules ^68^. We hypothesize that the interaction of TRAK1 or other adaptor proteins with molecular motors could differentially regulate axonal transport by fine-tuning the motility of molecular motors in microtubule regions coated by distinct cohesive islands.

Apart from providing additional anchoring points, TRAK1 enables processivity of KIF5B by the direct activation of the molecular motor in a kinesin-1 light chain-independent mechanism. In the absence of cargo, kinesin-1 is in an inactive conformation caused by the binding of its C-terminal cargo-binding domain to the N-terminal motor domain, which prevents the molecular motor from processively moving along microtubules ^32–34^. Binding of cargo to the tail of kinesin-1 heavy chain can activate processive motility of kinesin-1 (Coy et al., 1999; Jiang and Sheetz, 1995). The Drosophila Milton protein (TRAK ortholog) was found to interact directly with the kinesin-1 heavy chain without requiring kinesin-1 light chains ^39, 42^, and the TRAK2-binding site was mapped to the C-terminus of kinesin-1 KIF5A and KIF5C, respectively ^43, 44^, suggesting that TRAK could activate processive motility of kinesin-1. Here we provide direct evidence for this notion and show that kinesin-1 heavy chain KIF5B can recruit two TRAK1 dimers. To date only few proteins that activate kinesin-1 through a direct interaction with its heavy chain have been found, such as the Ran binding protein 2 (RanBP2) and Sunday Driver ^71, 72^. We found that TRAK1 belongs to this small group of proteins that directly activate kinesin-1 heavy chain KIF5B for processive motility (schematically shown in Figure S5).

By reconstituting mitochondrial transport *in vitro* using recombinant TRAK1 and KIF5B and isolated mitochondria, we showed that KIF5B and TRAK1 constitute the minimal transport complex that can drive directed mitochondrial motion along microtubules (schematically shown in Figure S5). The run length of these mitochondria increased in comparison to KIF5B and KIF5B-TRAK1, respectively, presumably due to the presence of multiple TRAK1-KIF5B transport complexes on a single mitochondrion, in line with previous studies on multi-motor transport ^22–25^. The additional anchoring of KIF5B by TRAK1 likely contributes to the increased run length of the mitochondria as predicted theoretically ^73^. The run length of mitochondria in axons of cultured hippocampal neurons is in the order of tens of micrometres ^20^ and the anterograde velocity can range from 0.1-0.8 μm/s ^41^ to 1 µm/s estimated in mice ^74^, comparable to values obtained in our experiments. Our *in vitro* reconstituted system, assembled using a minimal number of components, provides a powerful tool to test the mechanisms proposed to underpin the motion of mitochondria and to explore the regulatory roles of the individual components of the mitochondria trafficking machinery.

In summary, we demonstrate that TRAK1 anchors kinesin-1 to the microtubule surface, allowing for pausing instead of detaching when encountering obstacles, until either a blocked binding site is vacated or the motor site-steps. Tethering thus increases the processivity of kinesin-1, to enable microtubule-based transport in the crowded cytoplasmic environment. We propose tethering by auxiliary proteins as a general mechanism regulating molecular motors and other filament-associated proteins.

## Acknowledgments

We thank the Protein Facility of MPI-CBG Dresden and Yulia Bobrova for technical support. We acknowledge the financial support from GAUK (grant no. 250440 to V.H.), the Czech Science Foundation (grant no. 18-08304S to Z.L., 17-12496Y to M.B., 19-20553S to J.R. and 18-10832S to J.N.), the Australian Research Council Discovery (grant no. DP180103426 to J.N.), the Introduction of New Research Methods to BIOCEV (CZ.1.05/2.1.00/19.0390) project from the ERDF, the institutional support from the CAS (RVO: 86652036), the MS facility of CMS supported by MEYS CR (LM2015043) and the Imaging Methods Core Facility at BIOCEV, an institution supported by the Czech-BioImaging large RI projects (LM2015062 and CZ.02.1.01/0.0/0.0/16_013/0001775, funded by MEYS CR) for their support in obtaining imaging data presented in this paper.

## Authors contributions

Conceptualization, J.R., M.B., Z.L.; Methodology, V.H., J.R., C.B., M.B., Z.L.; Formal Analysis, V.H.; Investigation, V.H.; Resources, V.H., Z.N., L.G., S.D., J.N.; Writing -Original draft, V.H.; Review & Editing, V.H., J.N., S.D., J.R., M.B., Z.L.; Visualization, V.H.; Supervision, J.R., M.B., Z.L.; Funding Acquisition, V.H., J.R., J.N., M.B., Z.L.

## Declaration of interests

The authors declare no competing interests.

## Methods

### Production of recombinant proteins

Full-length mCherry-TRAK1 was cloned using Gateway Cloning. The cDNA encoding TRAK1 with an N-terminal mCherry-tag termed KIAA1042 (accession number Q9UPV9-1) was obtained from the Kazusa DNA Research Institute (Japan). The mCherry-TRAK1-encoding nucleotide sequences were PCR amplified using specifically designed primer pairs. For the generation of the gateway entry clone, the nucleotide sequence was inserted by means of a BP recombinant reaction (Invitrogen, Thermo Fisher Scientific, Carlsbad, CA, USA) according to the manufacturer’s protocol into a pDONR221 donor vector. The entry clone was verified by Sanger sequencing prior to transferring the entry clone into the destination vector. For the generation of the expression plasmid, an LR recombinant reaction was performed, generating a destination vector containing a TEV-cleavage site and a TwinStrep-FLAG-Halo-tag at the N-terminus of TRAK1. All TRAK1 constructs were expressed in HEK-293/T17 cells, grown in Free Style F17 medium (Gibco, Thermo Fisher Scientific, Inc., Waltham, MA, USA) supplemented with 0.1% Pluronic F-68 (Invitrogen, Thermo Fisher Scientific, Carlsbad, CA, USA) and 2 mM L-glutamine at 110 rpm under a humidified 5% CO2 atmosphere at 37 °C. 1 mg/ml linear polyethylene imine (Polysciences Inc., Warrington, PA, USA) and 0.7 mg of the expression plasmid were incubated in phosphate buffered saline (PBS) for 10 min prior to the addition to 350 ml cells at the concentration of 4 x 10^6^ cell/ml. Four hours post transfection, the cells were diluted two-fold in ExCell serum-free medium. Four days post transfection, the cells were harvested by centrifugation at 4 °C for 10 min at 500 x g. The cell pellet was resuspended in ice-cold lysis buffer (100 mM Tris-HCl, 10 mM NaCl, 5 mM KCl, 2 mM MgCl2, 10% glycerol, pH 8.0) supplemented with benzonase (1 U/ml; Merck, Darmstadt, Germany) and a protease inhibitor cocktail (cOmplete, EDTA free, Roche, Basel, Switzerland) and lysed by pulsed sonication for 5 min (20 s pulses with 24 W/min). Cell lysis was further assisted by the addition of Igepal-630 to the final concentration of 0.2% (v/v) during 20 min incubation on ice with occasional mixing. NaCl was added to the final concentration of 150 mM and the mixture was further incubated on ice for 20 min. Insoluble material was removed from by centrifugation at 9.000 x g for 15 min at 4 °C. The cell lysate was further cleared by a second centrifugation step at 30.000 x g for 30 min at 4 °C. The supernatant was loaded onto a StrepTactinXT column (IBA, Gottingen, Germany) equilibrated in the lysis buffer with 150 mM NaCl for affinity chromatography. After washing the column with wash buffer (100 mM Tris-HCl, 150 mM NaCl, 1 mM EDTA, pH 8), the protein was eluted by cleaving off the N-terminal tag with 1:20 (w/w) TEV protease in the wash buffer overnight at 4 °C. The next day, the eluted protein was collected, concentrated using an Amicon ultracentrifuge filter with a molecular weight cutoff of 100 kDa (Merck, Darmstadt, Germany) and loaded onto a Superose 6 10/300 GL column (GE Healthcare Bio-Sciences, Little Chalfont, UK) for further separation by size exclusion chromatography with 100 mM Tris-HCl pH 8.0, 150 mM NaCl, 2 mM MgCl2, 1 mM EDTA, 0.1% tween, 10% glycerol, 1mM DTT, 0.1 mM ATP as a mobile phase. The purified protein was concentrated using an Amicon ultracentrifuge filter and flash frozen in liquid nitrogen.

The mCherry-TRAK1 deletion mutant mCherry-TRAK1Δ was obtained by inserting a stop codon after amino acid 635 of the mCherry-TRAK1 encoding nucleotide sequence by means of a PCR-based mutagenesis (Agilent Technologies, QuikChange II Site-Directed Mutagenesis Kit) according to the manufacturers protocol. For the expression and purification of mCherry-TRAK1Δ the protocol as described above was followed. KIF5B constructs were obtained by PCR amplification of the amino acids 1-963 for full-length KIF5B and 1-905 for KIF5BΔ using primers containing AscI- and NotI-digestion sites flagging the particular KIF5B-encoding nucleotide sequences. After AscI-NotI-digestion of the inserts, they were ligated into an AscI-NotI-digested FlexiBAC destination vector containing a C-terminal fluorescent tag (GFP or mCherry) followed by a 3C PreScission protease cleavage site and a 6xHis-tag. All KIF5B constructs were expressed in SF9 insect cells using the opensource FlexiBAC baculovirus vector system for protein expression (Lemaitre et al., 2019). The insect cells were harvested after 4 days by centrifugation at 300 x g for 10 min at 4 °C in an Avanti J-26S ultracentrifuge (JLA-9.1000 rotor, Beckman Coulter, Brea, CA). The cell pellet was resuspended in 5 ml ice-cold PBS and stored at −80 °C for further use. For cell lysis, the insect cells were homogenized in 30 ml ice-cold His-Trap buffer (50 mM Na-phosphate buffer, pH 7.5, 5% glycerol, 300 mM KCl, 1 mM MgCl2, 0.1% tween-20, 10 mM BME, 0.1 mM ATP) supplemented with 30 mM imidazole, protease inhibitor cocktail and benzonase to the final concentration of 25 units/ml, and centrifuged at 45.000 x g for 60 min at 4 °C in the Avanti J-26S ultracentrifuge (JA-30.50Ti rotor, Beckman Coulter, Brea, CA). The cleared cell lysate was incubated for 2 h at 4 °C in a lysis buffer-equilibrated Ni-NTA column (HisPur Ni-NTA Superflow Agarose, Thermo Scientific, Thermo Fisher Scientific, Inc., Waltham, MA, USA) on a rotator for subsequent affinity chromatography via the C-terminal 6xHis-tag. The Ni-NTA column was washed with the wash buffer (His-Trap buffer supplemented with 60 mM imidazole) and the protein was eluted with the elution buffer (His-Trap buffer supplemented with 300 mM imidazole). The fractions containing the protein of interest were pooled, diluted 1:10 in the His-Trap buffer and the purification tag was cleaved overnight by 3C PreScisson protease. The solution was reloaded onto a Ni-NTA column to further separate the cleaved protein from the 6xHis-tag. The protein was concentrated using an Amicon ultracentrifuge filter and flash frozen in liquid nitrogen. The expression plasmid for the roadblock was an eGFP-labelled rigor binding kinesin-1 mutant from *Rattus norvegicus*, which contains the N-terminal 430 amino acids with a point mutation of amino acid 93 from threonine to asparagine and with a C-terminal eGFP- and 6xHis-tag ^31, 76^. The roadblock was expressed in *Escherichia coli* strain BL21(DE3) and purified via affinity chromatography using a Ni-NTA column as described. The final cleavage of the 6xHis-tag was omitted.

The human tau isoform htau441 with a C-terminal 6xHis- and mCherry-tag was expressed in SF9 insect cells and purified by affinity chromatography using the 6xHis-tag as described previously ^77^.

### Microtubules

Fluorescent microtubules were polymerized from 4 mg/ml porcine tubulin (80% unlabeled and 20% Alexa Fluor 647 NHS ester-labeled; Invitrogen, Thermo Fisher Scientific, Carlsbad, CA, USA) for 2 h at 37 °C in BRB80 (80 mM PIPES, 1 mM EGTA, 1 mM MgCl2, pH 6.9) supplemented with 1 mM MgCl2 and 1 mM GMPCPP (Jena Bioscience, Jena, Germany). The polymerized microtubules were centrifuged for 30 min at 18.000 x g in a Microfuge 18 Centrifuge (Beckman Coulter, Brea, CA) and the pellet was resuspended in BRB80 supplemented with 10 µM taxol (BRB80T).

### Preparation of cell extract for microscopy

Cell extracts of untransfected cells (native lysate) and cells transfected with DNA encoding TwinStrep-FLAG-Halo-mCherry-TRAK1 (TRAK1 lysate) or TwinStrep-FLAG-Halo-GFP (Halo lysate) were prepared from HEK-293/T17 cells. The cells were harvested by centrifugation for 5 min at 500 x g at 4 °C and the cell pellets were resuspended in 0.5 pellet volumes of lysis buffer (12 mM K-PIPES at pH 6.8, 1 mM MgCl2, 1 mM EGTA supplemented with 10 µg/ml cytochalasin D and a protease inhibitor cocktail), followed by pulsed sonication. Insoluble material was removed by centrifugation for 30 min at 20.000 x g at 4 °C. With the addition of 10 µg/ml cytochalasin D the polymerization of actin filaments was prevented. The cell lysate was used directly or was flash frozen in liquid nitrogen and stored at −80 °C for further use.

### Isolation of mitochondria

Murine mammary carcinoma cell line 4T1 stably expressing mitochondria-targeted eGFP was prepared by transfection with the pTagGFP2-mito vector (Evrogen, Moscow, Russia) using Lipofectamine 3000 (Thermo Scientific, Thermo Fisher Scientific, Inc., Waltham, MA, USA), followed by clonal selection with G418 (Sigma Aldrich, St. Louis, Missouri, USA). Cells were broken by a Balch-style homogenizer (Isobiotec, Heidelberg, Germany) set to 8 μm clearance and mitochondria were isolated using differential centrifugation as described before ^78, 79^. The mitochondrial pellet was washed 3x with isolation buffer (250 mM sucrose, 1 mM EDTA, 10 mM Tris/Mops pH 7.6) and resuspended in the same buffer, and the protein concentration was determined by a BCA assay (Thermo Scientific, Thermo Fisher Scientific, Inc., Waltham, MA, USA). Mitochondria were kept on ice and used within 4 hours in the microscopy experiments. Mitochondria isolated on three different days were analyzed by mass spectrometry to exclude the presence of co-purified TRAK or kinesin-1.

### *In vitro* molecule binding assay

Flow cells were prepared as described previously ^80^. Microtubules were immobilized in the flow cell by means of an anti-tubulin antibody (Sigma Aldrich, T7816, 10 µg/ml). Subsequently, the buffer was exchanged with motility buffer (BRB80 containing 10 µM taxol, 10 mM dithiothreitol, 20 mM D-glucose, 0.1% Tween-20, 0.5 mg/ml casein, 1 mM Mg-ATP, 0.22 mg/ml glucose oxidase and 20 µg/ml catalase). For experiments on bare microtubules, KIF5B constructs were diluted such that single molecules interacting with microtubules could be visualized (6 nM KIF5B-GFP, 0.2 nM KIF5BΔ-GFP or 1.3 nM KIF5BΔ-mCherry, respectively), or 175 nM TRAK1 constructs were flushed in the motility buffer directly into the flow cell, or a mixture of the two proteins was pre-incubated 10 min on ice prior to flushing into the flow cell for the imaging of KIF5B-TRAK1 complexes of various constructs. For decorating microtubules with roadblocks, 0.1 nM roadblocks in the motility buffer were flushed into the flow cell with immobilized microtubules. After an incubation for several seconds, unbound roadblocks were removed using the motility buffer. Subsequently, 2 nM KIF5BΔ-GFP or a pre-incubated complex of 2 nM KIF5BΔ-GFP and 100 nM mCherry-TRAK1, was flushed into the flow cell. For decorating microtubules with tau-mCherry islands, 3.5 nM tau-mCherry in the motility buffer was flushed into the flow cell containing immobilized microtubules. Tau-mCherry islands formed during the incubation time of up to five min, and unbound tau-mCherry was removed using the motility buffer. Subsequently, 2 nM KIF5BΔ-GFP in the absence or presence of 100 nM mCherry-TRAK1 in the motility buffer was added to the flow cell while keeping the tau-mCherry concentration in the solution constant. Experiments in cell lysates were performed as described previously ^30^: The cell lysates were incubated with an oxygen scavenger (20 mM glucose, 160 µg/ml glucose oxidase and 20 µg/ml catalase) for 20 min on ice followed by the addition of 2 mM MgATP and 0.2 nM KIF5BΔ-GFP prior to flushing into the flow cell. Mitochondrial transport was reconstituted by pre-incubating 2 nM unlabeled KIF5BΔ, 100 nM mCherry-TRAK1 and 10 µg mitochondria-GFP on ice prior to flushing the mixture into the flow cell with immobilized microtubules. All experiments were performed at room temperature.

### Total internal reflection fluorescence microscopy and imaging acquisition

For fluorescence imaging, the total internal reflection fluorescence (TIRF) mode of an inverted widefield fluorescence microscope (Nikon Eclipse Ti-E; Nikon, Tokyo, Japan) equipped with a motorized XY stage and a perfect focus system was used together with a 60x oil immersion objective (Nikon CFI Apo TIRF 60x Oil, NA 1.49, WD 0.12 mm), a 2.5x relay lens in front of an electron-multiplied charge-coupled device camera (iXon Ultra DU-888; Andor, Belfast, Northern Ireland) and, if necessary, an additional 1.5x magnifying tube lens. Alexa647-labelled microtubules, mCherry- and GFP-labelled proteins and GFP-labelled mitochondria were visualized by the sequential switching between a Cy5 filter (632-652, 669-741), TRITC filter (556-566, 593-668) and FITC filter (483-493, 500-550) or by using a Quad Band Set filter (405/488/561/640). Images were acquired for one to two minutes with 200 ms exposure time and 300 gain multiplier using NIS-Elements Advanced Research software v5.02 (Laboratory Imaging). Experiments were performed over several months, each experiment presented was repeated at least on three individual days. No data were excluded from the study.

### Fluorescence image analysis

Image analysis for estimating the motility parameters (interaction time, run length and velocity in Figures 3-5, S3a, S4c, S4d) of fluorescently labelled KIF5B and TRAK1 constructs was performed by tracking the movement of the respective molecules with the high-precision tracking software FIESTA ^80^. All trajectories were double-checked by eye to avoid computer misinterpretations. The following trajectories were not analyzed: stationary molecules not exhibiting processive movements, clustering molecules, molecules passing crossing microtubules and stationary molecules accumulating at the microtubule plus-end. For a Kaplan-Meier estimation, the following trajectories were denoted as censored: trajectories reaching the end of a microtubule, trajectories starting in the first or ending in the last imaging frame, trajectories starting or ending at the edge of the field of view. Further trajectory evaluation was performed in MatLab (The MathWorks, Natick, MA, USA). The interaction time was determined as the time difference between the beginning and the end of the trajectory and the run length as the distance along the path of the trajectory. Kaplan-Meier estimations were evaluated using the MatLab build-in empirical cumulative distribution function (ecdf), which computes the 95% confidence interval using the Greenwood’s formula. Calculated velocities represent the average velocity of a molecules from the beginning to the end of their interaction with a microtubule (Figure 3 and 5). Frame-to-frame velocities (Figure S3a) were computed by extracting the position of a molecule in each frame and calculating the velocities between two consecutive frames. The run length and interaction time of molecules within tau-mCherry islands were determined by manually measuring the time and position of the molecule when entering the tau island and when leaving the tau-mCherry island by either passing the whole island or dissociating from the microtubule, respectively, using the image processing software Fiji ^81^. Molecules that did not enter tau-mCherry islands were not included in the evaluation. The run length and velocities of mitochondria (Figure 5) were manually measured by determining the coordinates of the beginning and the end of the movements of mitochondria using the Fiji software. Histograms of the integrated fluorescence intensity distributions of the GFP- and mCherry-signals of KIF5BΔ and TRAK1, respectively, (Figure 2c, 2d) were determined by measuring the integrated intensity of the first four images of a movie taken of a molecule after binding to the microtubule in a square of 9x9 pixels and subtracting the integrated intensity of the background in a square of 9x9 pixels adjacent to the microtubule at the same time frames using the Fiji software. Values higher than 6000 a.u., representing less then 18% of the fluorescent particles, likely caused by unspecific clustering of molecules, were disregarded. Time traces of fluorescent intensities (Figure S2) were determined by a line scan along the trace of the molecule of interest in the respective kymograph.

## Contact for resource sharing

Further information and requests for reagents can be directed to and will be fulfilled by Marcus Braun and Zdenek Lansky.

---

**Figure S1.**
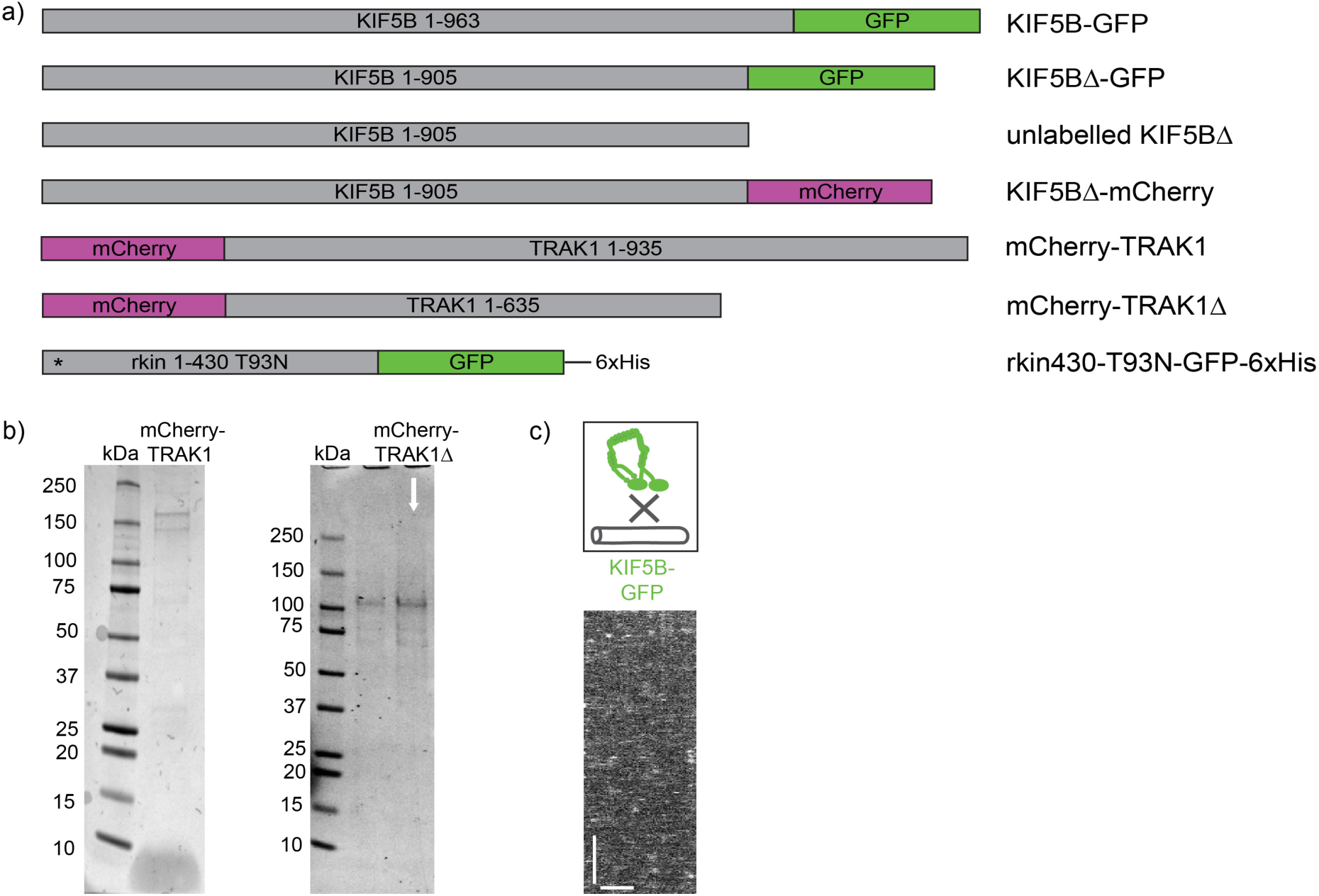
Related to Figure 1-5. **a)** List of constructs used in this study. The asterisk indicates the position of the T93N point mutation of the rigor binding kinesin-1 mutant rkin430. **b)** SDS-gels of purified mCherry-TRAK1 (left) and mCherry-TRAK1Δ (right). **c)** Schematic illustration and kymograph of auto-inhibited KIF5B-GFP not interacting with the microtubule (compare with the activated KIF5B-GFP in Figure 1b). Scale bars: 2 µm, 10 s.

**Figure S2.**
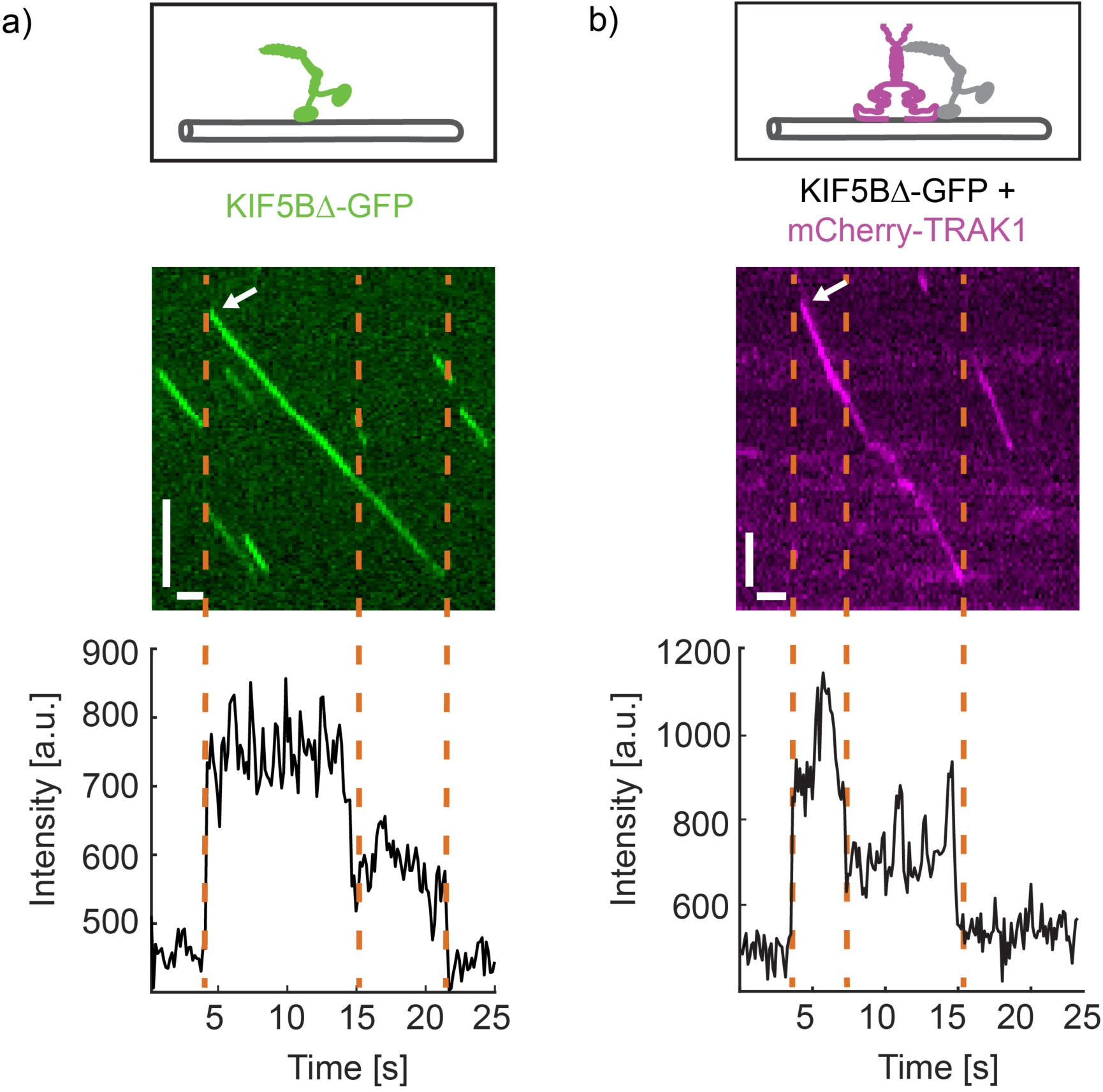
Related to Figure 2. **a)** Schematic illustration, kymograph and intensity time trace of KIF5BΔ-GFP moving processively along a microtubule. The intensity time trace of the KIF5BΔ-GFP molecule shown in the kymograph (white arrow) exhibits two bleaching steps (indicated by the dashed lines) of the two GFP-fluorophores of the dimeric molecular motor. **b)** Schematic illustration, kymograph and intensity time trace of mCherry-TRAK1 transported by KIF5BΔ-GFP along a microtubule. The intensity time trace of the mCherry-TRAK1 molecule shown in the kymograph (white arrow) exhibits two bleaching steps (indicated by the dashed lines) of the two mCherry-fluorophores of mCherry-TRAK1, indicating its dimeric form. Scale bars: 2 s, 2 µm. Intensity time traces were obtained by a line scan along the traces of the molecules within the kymographs (Methods).

**Figure S3.**
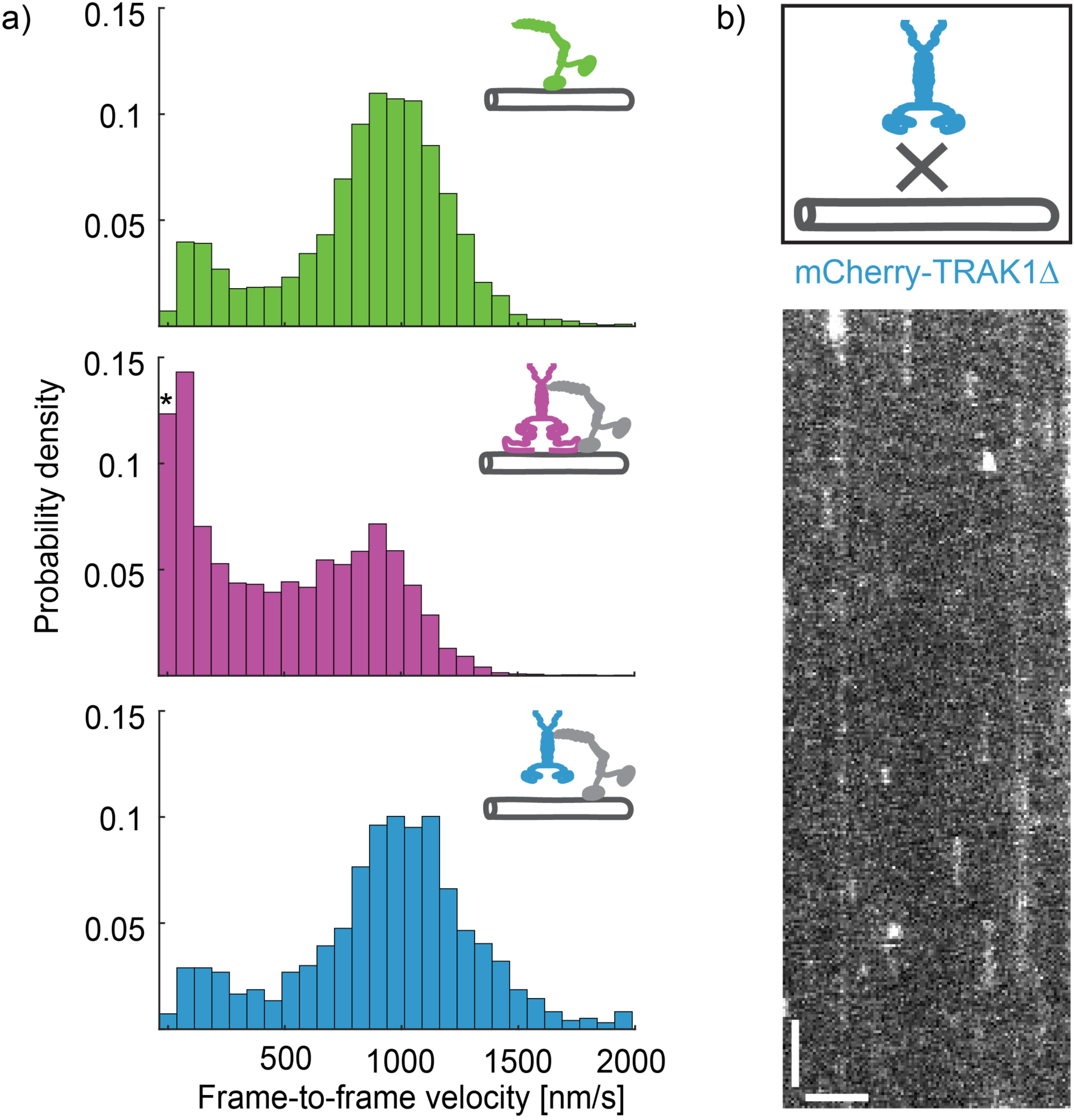
Related to Figure 3. **a)** Histograms of frame-to-frame velocities of KIF5BΔ-GFP (green, top), KIF5BΔ-GFP in the presence of mCherry-TRAK1 (magenta, middle) and KIF5BΔ-GFP in the presence of mCherry-TRAK1Δ (cyan, bottom). The median frame-to-frame velocity of KIF5BΔ-GFP decreased in the presence of mCherry-TRAK1 from 909 nm/s (lower, upper interquartile limit 282 nm/s, 1085 nm/s, n = 534 molecules) to 458 nm/s (lower, upper interquartile limit 98 nm/s, 842 nm/s, n = 498 molecules). The population around zero (indicated by an asterisk) represents a high number of transient pauses. In the presence of mCherry-TRAK1Δ the median frame-to-frame velocity of KIF5BΔ-GFP barely decreased (972 nm/s, lower, upper interquartile limit 282 nm/s, 1085 nm/s, n = 83 molecules). **b**) Schematic representation and kymograph of mCherry-TRAK1Δ, a TRAK1-construct lacking the microtubule-binding domain, not interacting with microtubules (compare to mCherry-TRAK1 diffusing along a microtubule in Figure 1a). Scale bars: 2 µm, 5 s.

**Figure S4.**
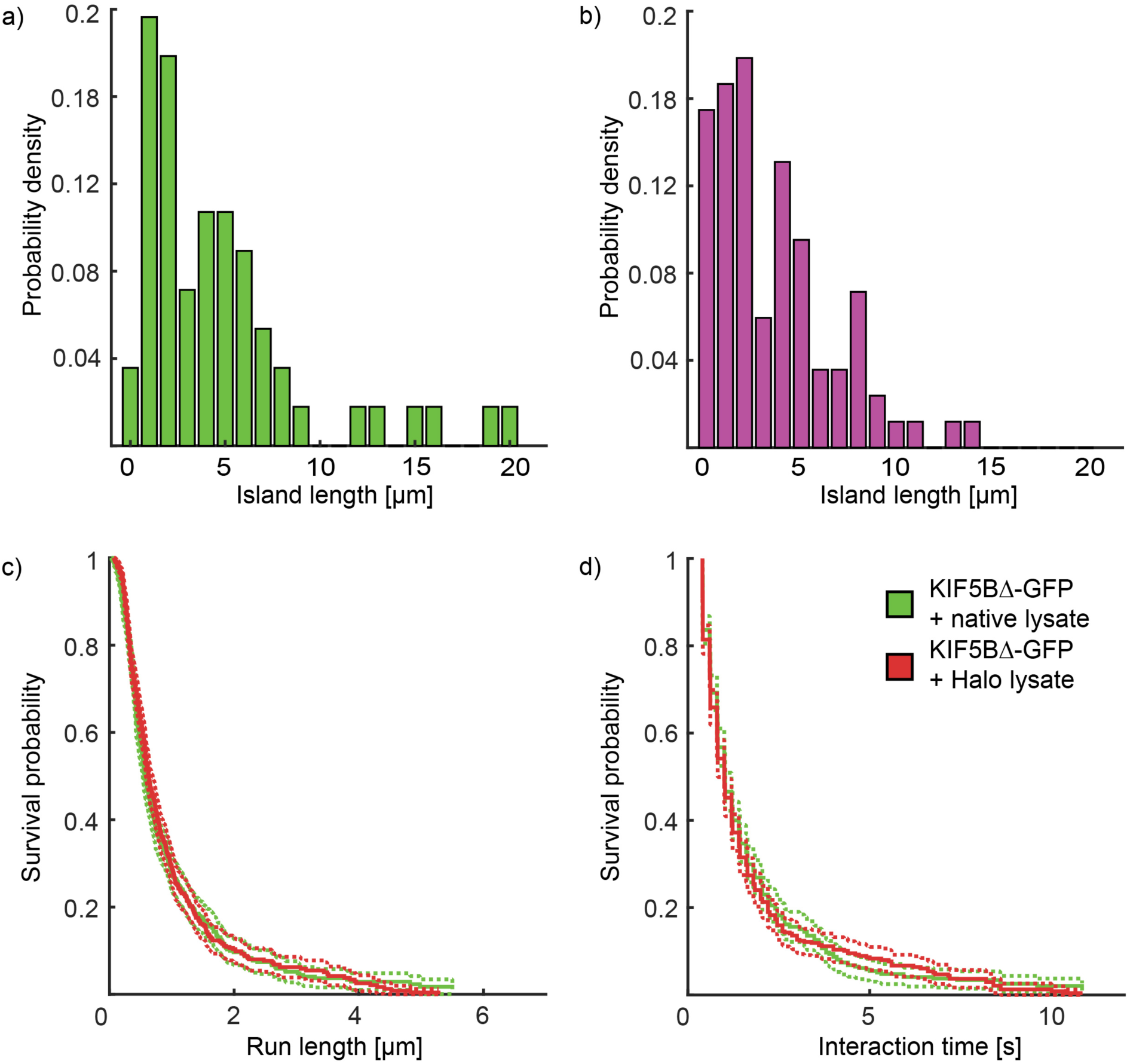
Related to Figure 4. Histograms of the tau island sizes in experiments with **a)** KIF5BΔ-GFP (left, 4.75 ± 4.8 µm, mean ± standard deviation) and **b)** KIF5BΔ-GFP-mCherry-TRAK1 complexes (right, 3.54 ± 3.12 µm, mean ± standard deviation). **c), d)** Survival probability plots showing a similar median run length and median interaction time of KIF5BΔ-GFP in native (green) and Halo-expressing (red) cell lysate.

**Figure S5.**
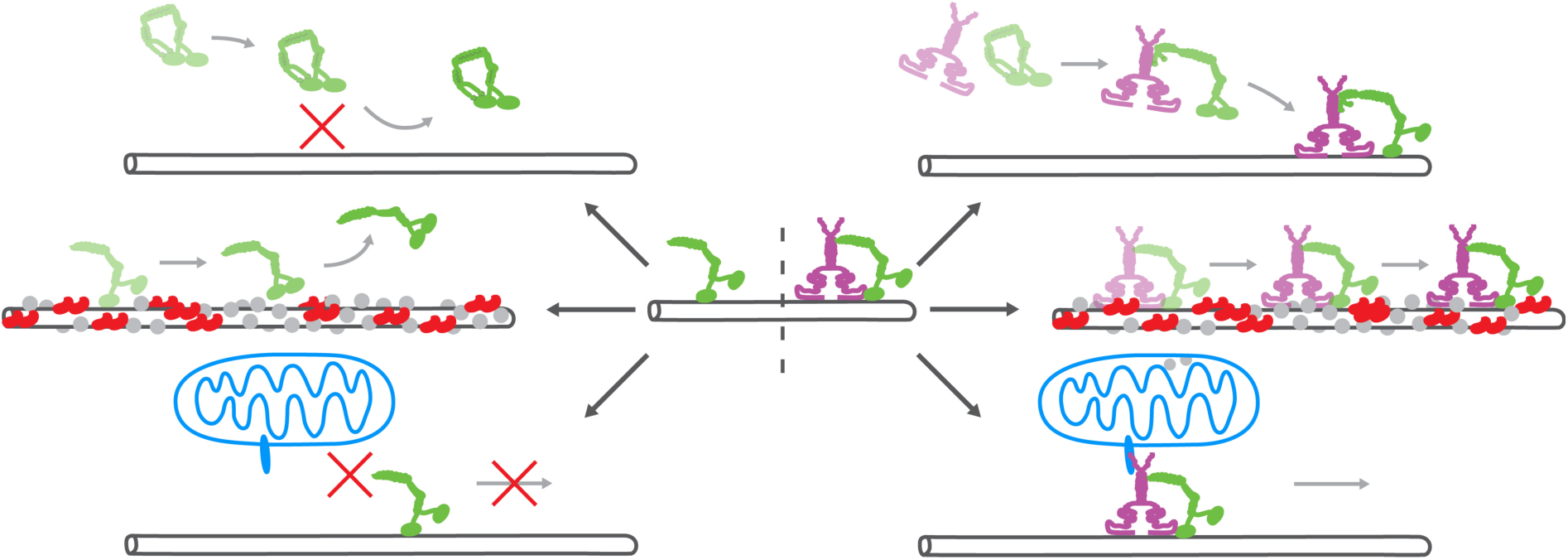
Related to Figure 1, 4 and 5. Overview of the functions of TRAK1. Top: TRAK1 activates auto-inhibited KIF5B, enabling its processive movement along microtubules. Middle: TRAK1 increases the processivity of KIF5B in crowded environments. Bottom: TRAK1 enables KIF5B-based transport of isolated mitochondria along microtubules *in vitro*.

